# ATR promotes mTORC1 activity via *de novo* cholesterol synthesis

**DOI:** 10.1101/2023.10.27.564195

**Authors:** Naveen Kumar Tangudu, Alexandra N. Grumet, Richard Fang, Raquel Buj, Aidan R. Cole, Apoorva Uboveja, Amandine Amalric, Baixue Yang, Zhentai Huang, Cassandra Happe, Mai Sun, Stacy L. Gelhaus, Matthew L. MacDonald, Nadine Hempel, Nathaniel W. Snyder, Katarzyna M. Kedziora, Alexander J. Valvezan, Katherine M. Aird

## Abstract

DNA damage and cellular metabolism exhibit a complex interplay characterized by bidirectional feedback mechanisms. Key mediators of the DNA damage response and cellular metabolic regulation include Ataxia Telangiectasia and Rad3-related protein (ATR) and the mechanistic Target of Rapamycin Complex 1 (mTORC1), respectively. Previous studies have established ATR as a regulatory upstream factor of mTORC1 during replication stress; however, the precise mechanisms by which mTORC1 is activated in this context remain poorly defined. Additionally, the activity of this signaling axis in unperturbed cells has not been extensively investigated. Here, we demonstrate that ATR promotes mTORC1 activity across various cellular models under basal conditions. This effect is particularly enhanced in cells following the loss of p16, which we have previously associated with hyperactivation of mTORC1 signaling and here found have increased ATR activity. Mechanistically, we found that ATR promotes *de novo* cholesterol synthesis and mTORC1 activation through the upregulation of lanosterol synthase (LSS), independently of both CHK1 and the TSC complex. Furthermore, the attenuation of mTORC1 activity resulting from ATR inhibition was rescued by supplementation with lanosterol or cholesterol in multiple cellular contexts. This restoration corresponded with enhanced localization of mTOR to the lysosome. Collectively, our findings demonstrate a novel connection linking ATR and mTORC1 signaling through the modulation of cholesterol metabolism.

## INTRODUCTION

The DNA damage response (DDR) plays an important role in maintaining genome stability (Jeggo et al., 2016). This is particularly apparent in cancer where cells hijack DDR by upregulation of the key DDR proteins Ataxia Telangiectasia and Rad3-related protein (ATR) and/or Ataxia Telangiectasia Mutated (ATM) (Weber and Ryan, 2015). While activation of these pathways can induce cell cycle arrest to allow for DNA damage repair, cancers often have other cell cycle or DDR alterations that allow for continued proliferation even in the presence of activated ATR/ATM (Maréchal and Zou, 2013). Indeed, many cancers are often reliant on these proteins for continued proliferation, and ATR and ATM inhibitors are in clinical trials for a variety of malignancies (Priya et al., 2023). Additionally, recent work has demonstrated a critical role for ATR beyond the DDR in normal proliferation (Menolfi et al., 2023; Sugitani et al., 2022), suggesting that ATR activity under basal conditions plays an active role in cell signaling. Pathways downstream of ATR beyond cell cycle control and DNA repair remain unclear.

DDR and metabolism are bidirectionally linked (Cucchi et al., 2021; Turgeon et al., 2018). We and others have shown that ATR and ATM have roles in metabolism (Aird et al., 2015; Chen et al., 2020; Dahl and Aird, 2017; Diehl et al., 2022; Huang et al., 2023), and metabolism can also influence DNA repair by regulating repair components and providing substrates necessary for DNA repair (Chatzidoukaki et al., 2020; Cucchi et al., 2021; Uboveja and Aird, 2024). It is therefore unsurprising that studies have previously identified a bidirectional relationship between both ATR and ATM to the master nutrient sensor and metabolic regulator mechanistic Target of Rapamycin Complex 1 (mTORC1) (Alexander et al., 2010; Danesh Pazhooh et al., 2021; Lamm et al., 2020; Ma et al., 2018), although the mechanisms linking these pathways have not yet been fully explored. We previously published that loss of the tumor suppressor p16 leads to mTORC1 hyperactivation in both normal and cancer cells (Buj et al., 2019). The mechanism underlying how p16 loss enhances mTORC1 activity in our model is not yet understood. We also found that p16 loss increases DNA damage (Tangudu et al., 2024). Whether the DDR and mTORC1 are linked downstream of p16 loss, or in other contexts, and the mechanism underlying this remains unclear.

mTORC1 is a master regulator of anabolic cell growth and proliferation that links upstream nutrient, growth factor, and energy signals to downstream metabolic pathways (Valvezan and Manning, 2019). mTORC1 activity is regulated through control of its spatial localization to the cytosolic surface of the lysosome. Sufficient levels of intracellular nutrients, including cholesterol, amino acids, and glucose, promote mTORC1 localization to the lysosome where it can be activated by the small GTPase Rheb (Castellano et al., 2017; Saxton and Sabatini, 2017; Shin et al., 2022). mTORC1 activation also requires inhibition of its upstream negative regulator, the TSC complex, which also resides at the lysosome. This often occurs downstream of growth factor signaling pathways including PI3K/Akt, MEK/Erk, and others, which cause the TSC complex to dissociate from the lysosome allowing mTORC1 activation (Ilagan and Manning, 2016; Menon et al., 2014). While ATR and ATM have been linked to mTORC1 activity (Alexander et al., 2010; Danesh Pazhooh et al., 2021; Lamm et al., 2020; Ma et al., 2018), whether and how these pathways affect mTORC1 lysosomal localization and activation is unknown.

Cholesterol is a key component of the plasma membrane and also plays important roles in signaling (Liu et al., 2023). The *de novo* cholesterol synthesis pathway is a multi-step process starting with acetyl-CoA and progressing through mevalonate, squalene, and lanosterol, culminating in cholesterol (Shi et al., 2022). Prior work has shown that cholesterol activates mTORC1 via LYCHOS at lysosomes (Castellano et al., 2017; Davis et al., 2021; Lim et al., 2019; Shin et al., 2022), and cholesterol has also been linked to genome instability and the DDR (Liu et al., 2023). Whether the DDR and mTORC1 are linked through cholesterol has not been previously studied.

Here we observed that knockdown or inhibition of ATR decreases mTORC1 activity in multiple cell models to a greater extent than suppression of ATM, and that the ATR-mTORC1 signaling axis was especially apparently in p16 knockdown cells, which have mTORC1 hyperactivation (Buj et al., 2019). Using an unbiased proteomics approach cross-compared with publicly available datasets of potential upstream regulators of mTORC1, we identified the *de novo* cholesterol synthesis enzyme lanosterol synthase (LSS) as a mediator between ATR and mTORC1 activation in p16 knockdown cells. Indeed, p16 knockdown cells showed increased cholesterol abundance in an ATR-dependent manner. Moreover, in multiple cellular models, suppression of mTORC1 activity by ATR inhibition was rescued by exogenous cholesterol or lanosterol. Suppression of ATR also decreased mTOR localization at the lysosome, which was rescued by cholesterol. Together, our data demonstrate an unexpected link between ATR and mTORC1 via cholesterol synthesis.

## RESULTS

### ATR and ATM suppression decrease mTORC1 activity in unperturbed cells independently of TSC2

Prior work has placed ATR and ATM upstream of mTORC1 under conditions of replication stress and/or DNA damage (Alexander et al., 2010; Danesh Pazhooh et al., 2021; Lamm et al., 2020; Ma et al., 2018). To explore whether ATR/ATM and mTORC1 are linked under unperturbed basal conditions, we inhibited or knocked down ATR and ATM in multiple cell models and assessed phosphorylation of the direct mTORC1 substrate S6K (pS6K) as a well-established readout of mTORC1 activity. Suppression of ATR or ATM, but not CHK1 or CHK2, decreased pS6K (**Fig. 1A-C and S1**). Surprisingly the dual CHK1/CHK2 inhibitor AZD7762 strongly reduced pS6K; however, this appears to be an off-target effect as it was not reproduced with another dual

**Figure 1.**
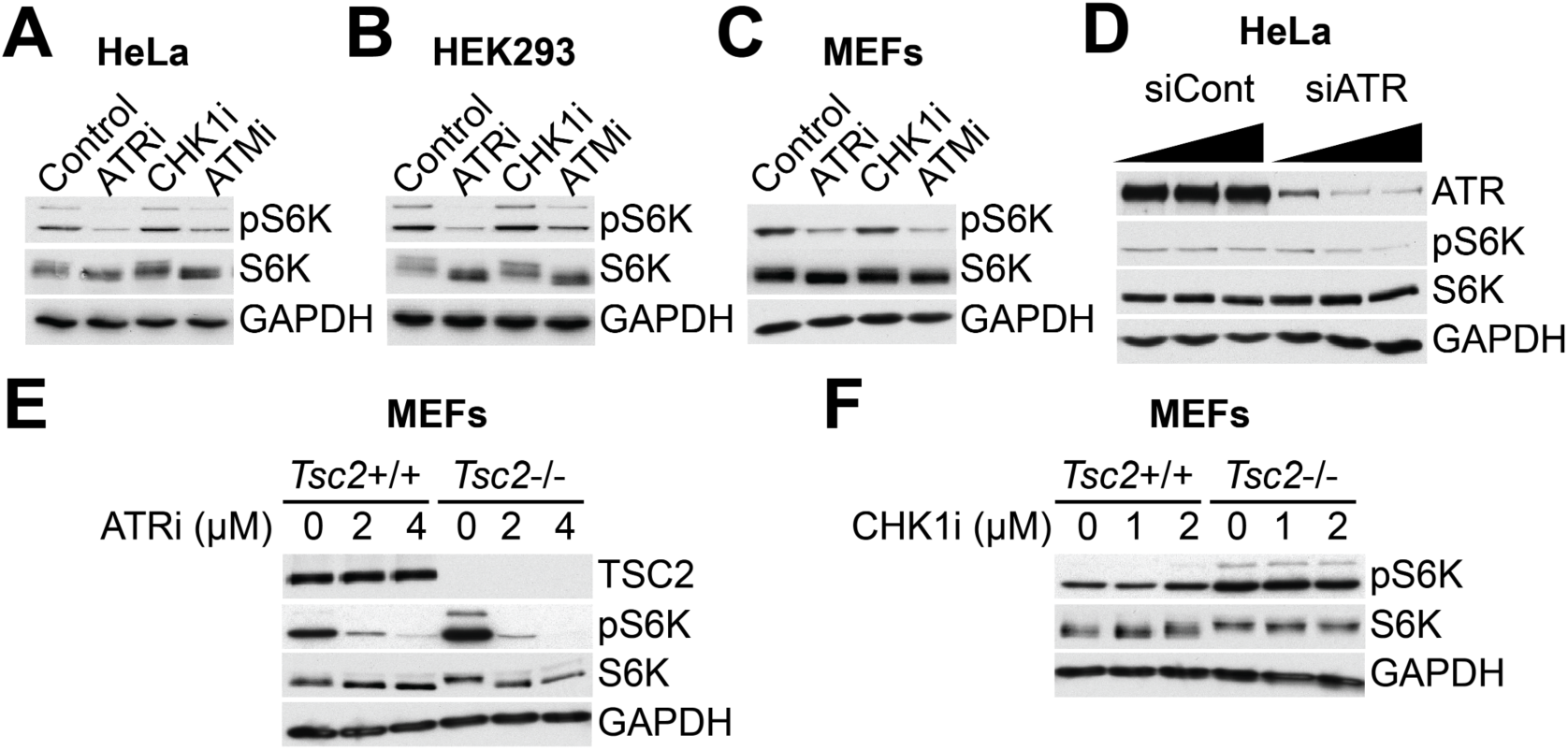
ATR or ATM inhibition and knockdown suppress mTORC1 activity independent of TSC2. **(A)** HeLa cells, **(B)** HEK293 cells, and **(C)** MEFs (mouse embryonic fibroblasts) were treated for 3h with 2μM AZD6738 ATR inhibitor (ATRi), 2μM LY2603618 CHK1 inhibitor (CHK1i), or 5μM KU55933 ATM inhibitor (ATMi), and the indicated proteins were assessed by western blotting. GAPDH was used as a loading control. **(D)** HeLa cells were transfected with siRNA against ATR or with a non-targeting siRNA as control, and the indicated proteins were assessed by western blotting. GAPDH was used as a loading control. **(E-F)** MEFs with TSC2 knockout were treated for 3h with the indicated doses of AZD6738 ATR inhibitor (ATRi) **(E)** or LY2603618 CHK1 inhibitor (CHK1i) **(F)**, and the indicated proteins were assessed by western blotting. GAPDH was used as a loading control. All western blots are representative data from at least 3 independent experiments.

CHK1/CHK2 inhibitor Prexasertib, or with combined CHK1 inhibitor plus CHK2 knockdown (**Fig. S1**). Importantly our ATR siRNA/shRNA knockdown approaches are key to demonstrating that the ATR inhibitor effects are not simply due to off-target effects on mTOR itself, which is a known effect of previous generation ATR inhibitors (Priya et al., 2023) **(Fig. 1D)**. The TSC complex is a potent negative regulator of mTORC1 that integrates many upstream signals to control mTORC1 activity (Huang and Manning, 2008). Loss of the essential TSC complex component TSC2 did not affect ATR inhibitor-mediated reduction of pS6K, demonstrating the effect is independent of the TSC complex **(Fig. 1E)**. CHK1 inhibition did not affect S6K phosphorylation regardless of TSC2 status **(Fig. 1F)**. Together, these data demonstrate a link between the DDR, and in particular ATR, and mTORC1 signaling in unperturbed cells and that is independent of the TSC complex and CHK1.

### ATR signaling downstream of p16 loss promotes mTORC1 activity

We found that suppressing ATR and ATM decreased mTORC1 activation in unperturbed cells (**Fig. 1**). Next, we aimed to test whether this pathway is intact in cells with combined high DDR and mTORC1 hyperactivation. Cells with knockdown of p16 have both increased DNA damage (Tangudu et al., 2024) and hyperactive mTORC1 signaling (Buj et al., 2019). Thus, we aimed to determine whether p16 knockdown cells increase mTORC1 activity via ATR and/or ATM using a previously validated shRNA against p16 (Buj et al., 2019). Consistent with the increase in DNA damage observed in p16 knockdown cells (Tangudu et al., 2024), we found that these cells have increased phosphorylation of ATR/ATM substrates pCHK1 and pCHK2 (**Fig. S2A**). Additionally, using melanoma patient samples from TCGA, we found that homozygous deletion of *CDKN2A*, the gene encoding p16, correlated with increased pCHK1 along with other proteins involved in DNA damage response and repair (**Fig. S2B**). Together, these data demonstrate that loss of p16 expression promotes ATR/ATM pathway activation, and this model is therefore useful towards understanding the mechanistic link between ATR/ATM and mTORC1. Knockdown or inhibition of ATR in p16 knockdown cells robustly decreased pS6K (**Fig. 2A-B**), while ATM knockdown only modestly decreased pS6K, and the ATM inhibitor KU60019 had no effect (**Fig. 2C-D)**. This was not due to a lack of ATM inhibitor efficacy as we noted robust downregulation of pCHK2 (**Fig. 2D**). Knockdown of p16 also strongly sensitized mTORC1 activity to ATR inhibition in HeLa cells (**Fig. S2C)**, although we do note that p16 knockdown itself in these cells did not robustly increase pS6K, which may be due to differential genetics and signaling between these cells and the cancer cell lines we have previously published (Buj et al., 2019). Similar to unperturbed cells **(Fig. 1A-C**), knockdown of CHK1 in p16 knockdown cells did not robustly decrease pS6K (**Fig. S2D**), demonstrating that ATR promotion of mTORC1 activity is independent of this key downstream ATR substrate. Finally, we aimed to confirm the directionality of the effect as prior work has placed mTORC1 upstream of ATR (Danesh Pazhooh et al., 2021; Ma et al., 2018). Inhibition of mTORC1 signaling using Torin1 did not decrease pCHK1 in p16 knockdown cells (**Fig. 2E**). Together, these results demonstrate that ATR is upstream of mTORC1 and promotes its activation in p16 knockdown cells.

**Figure 2.**
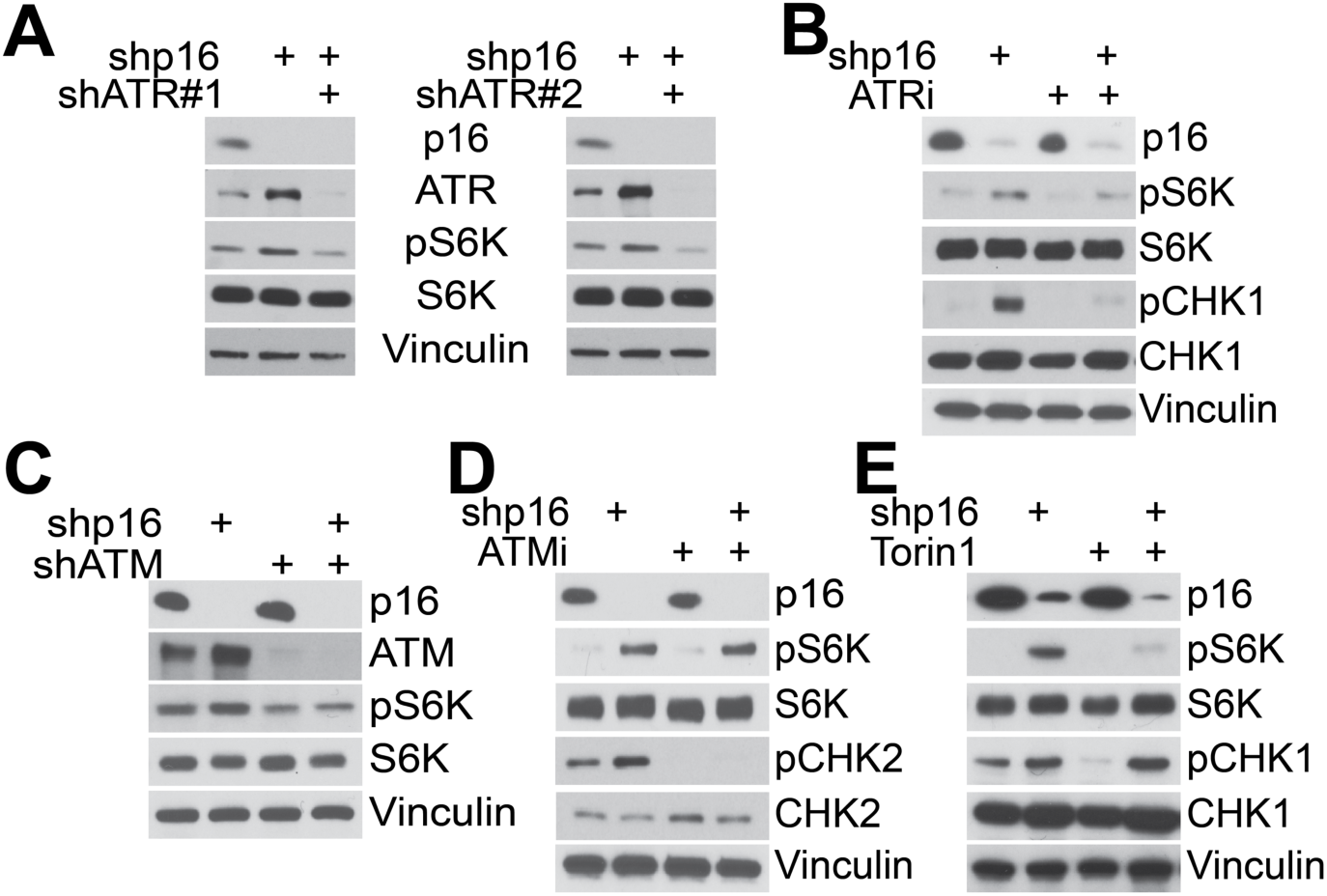
ATR affects mTORC1 activity in p16 knockdown cells. SKMEL28 cells were transduced with a lentivirus expressing a short hairpin RNA (shRNA) targeting GFP or p16 (shp16). **(A)** shp16 cells were transduced with a lentivirus expressing a shRNA targeting GFP or two independent shRNAs targeting ATR (shATR #1 and #2), and the indicated proteins were assessed by western blotting. Vinculin was used as a loading control. **(B)** Cells were treated for 30min with 63nM AZD6738 ATR inhibitor (ATRi), and the indicated proteins were assessed by western blotting. Vinculin was used as a loading control. **(C)** shp16 cells were transduced with a lentivirus expressing a shRNA targeting GFP or ATM (shATM), and the indicated proteins were assessed by western blotting. Vinculin was used as a loading control. Cells were treated for 30min with 313nM KU60019 ATM inhibitor (ATMi), and the indicated proteins were assessed by western blotting. Vinculin was used as a loading control. **(E)** Cells were treated for 10min with 250nM of the mTORC1 inhibitor Torin1, and the indicated proteins were assessed by western blotting. Vinculin was used as a loading control. All western blots are representative data from 3 independent experiments.

### ATR increases lanosterol synthase (LSS) and intracellular cholesterol to promote mTORC1 activity

We next aimed to understand the mechanism that couples ATR to mTORC1 activity in p16 knockdown cells. We cross-compared proteomics data of p16 knockdown cells with knockdown of ATR (**Table S1**) to a publicly available dataset of potential mTORC1 regulators (Condon et al., 2021) (**Fig. 3A**). This resulted in 9 potential upstream mTORC1 regulators that were specific to p16 knockdown cells (**Fig. 3B**), which have high ATR-mediated mTORC1 activity (**Fig. 2**). The protein that was most significantly downregulated by ATR knockdown was lanosterol synthase (LSS) (**Fig. 3B-C**). LSS is an enzyme that forms the four-ring structure of cholesterol during the conversion of (*S*)-2,3-epoxysqualene to lanosterol in the *de novo* cholesterol synthesis pathway (**Fig. 3D**). We validated increased LSS in two different cell line models with p16 knockdown, which was subsequently reduced by ATR knockdown (**Fig. 3E and S3A**). Consistent with ATR promoting LSS expression in p16 knockdown cells, we observed increased intracellular cholesterol levels using filipin staining that was subsequently decreased by ATR knockdown in both cell lines (**Fig. 3F-G and S3B-C**). Together, these data demonstrate that ATR regulates LSS expression and cholesterol levels in p16 knockdown cells.

**Figure 3.**
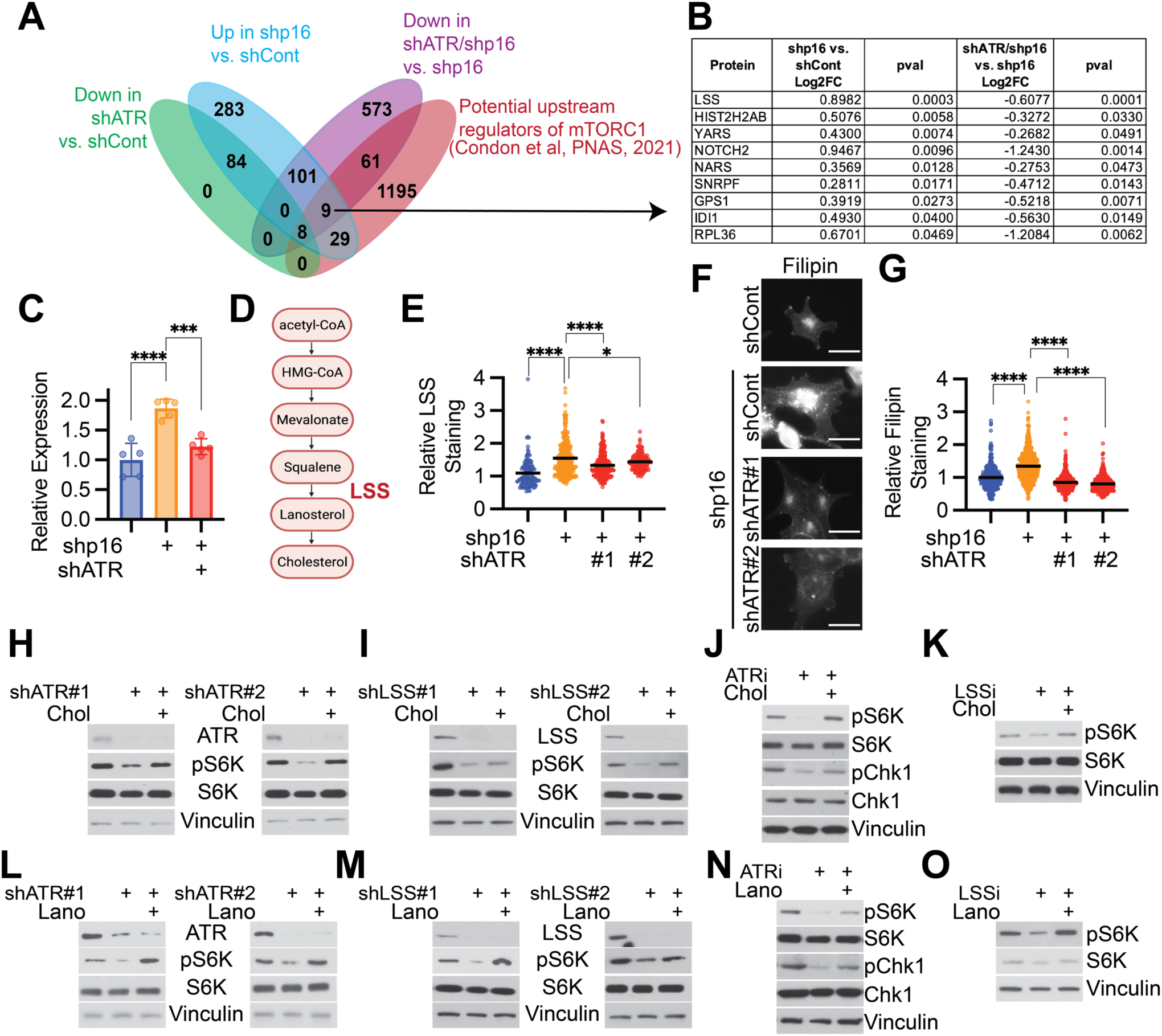
ATR decreases lanosterol synthase expression and cholesterol to regulate mTORC1 activity. (A-G) SKMEL28 cells were transduced with lentivirus expressing shRNA targeting GFP (shCont) or p16 (shp16) with or without lentivirus expressing shRNA targeting ATR (shATR #1 and #2). **(A)** Venn diagram comparing proteomics data with publicly available dataset of mTORC1 regulators. **(B)** Nine hits were identified in the cross-comparison from **(A)**. **(C)** Relative LSS protein abundance in the indicated groups was determined by mass spectrometry. Graph represents mean ± SD. **(D)** Simplified pathway of *de novo* cholesterol synthesis and LSS. **(E)** LSS expression was assessed by immunofluorescence staining and quantified. Graph represents individual normalized values and mean. **(F-G)** Cholesterol abundance was assessed by filipin staining **(F)** and quantified **(G)**. Scale bar = 20μm. Graph represents individual normalized values and mean. **(H-O)** shp16 SKMEL28 cells were transduced with lentivirus expressing shRNA targeting ATR (shATR #1 and #2) **(H and L)** or targeting LSS (shLSS #1 and #2) **(I and M).** SKMEL28 cells were treated with 63nM AZD6738 ATR inhibitor (ATRi) **(J and N)** or 2.5μM LSS inhibitor (LSSi) Ro 48-8071 for 2h **(K and O)**. The indicated proteins were assessed by western blotting. Vinculin was used as a loading control. **(H-K)** Cells were supplemented with 50μM cholesterol. **(L-O)** Cells were supplemented with 50μM lanosterol. Representative data from one of three independent experiments is shown. One-way ANOVA. ***p<0.005, ****p<0.001

Prior work found that cholesterol promotes mTORC1 activation (Castellano et al., 2017; Davis et al., 2021; Lim et al., 2019; Shin et al., 2022). Consistent with the ATR-LSS axis promoting mTORC1 activity via cholesterol in p16 knockdown cells, knockdown or inhibition of either ATR or LSS decreased mTORC1 activity, and these effects were rescued by supplementation with either lanosterol or cholesterol (**Fig. 3H-O and S3D-G**). The decrease in mTORC1 activity upon ATR inhibition in HeLa cells was also rescued by supplementation with cholesterol (**Fig. S3H**). Together, our data demonstrate that ATR upregulates LSS, which promotes mTORC1 activity through cholesterol.

### ATR promotes mTOR localization to lysosomes via cholesterol

mTOR localization to the lysosome in response to nutrients facilitates its activation (Castellano et al., 2017; Kim et al., 2008; Sancak et al., 2008). Thus, we aimed to determine whether ATR promotes mTOR lysosomal localization through its effects on cholesterol. Indeed, knockdown or inhibition of ATR decreased mTOR localization to the lysosome, and this effect was rescued by exogenous cholesterol supplementation (**Fig. 4A and S4A**). Consistent with ATR promoting mTORC1 activation independent of the TSC complex (**Fig. 1E**), ATR knockdown or inhibition did not increase TSC2 lysosomal localization (**Fig. 4B and S4B**). Surprisingly, ATR knockdown slightly reduced TSC2 lysosomal localization in p16 knockdown SKMEL28 cells, although this did not occur with ATR inhibition in HeLa cells and was not affected by cholesterol (**Fig. 4B and S4B**). Taken together with results in Figure 3, these data put forth a model in which ATR promotes LSS expression and increased intracellular cholesterol levels, which activate mTORC1 by promoting its lysosomal localization (**Fig. 4C**).

**Figure 4.**
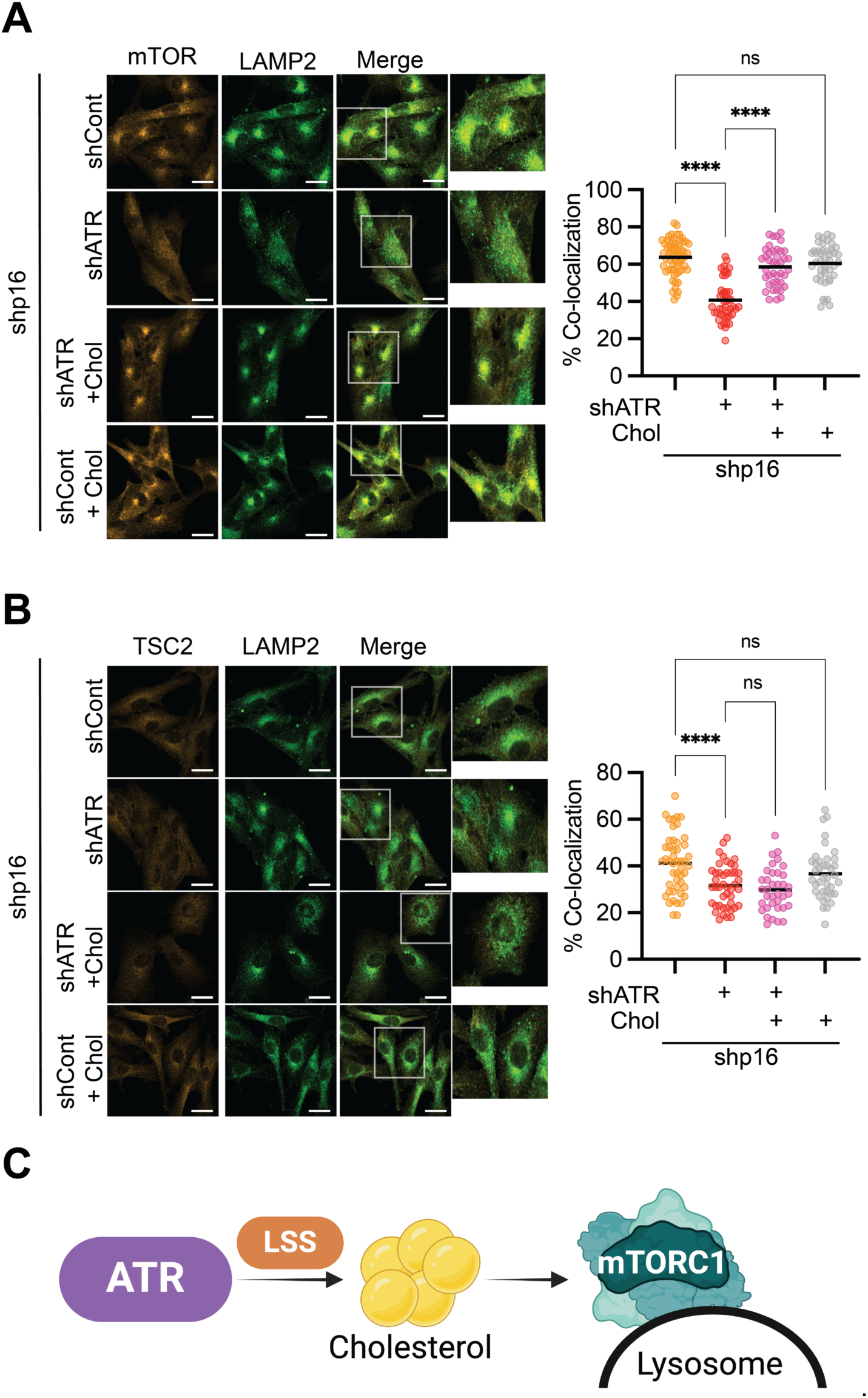
ATR promotes mTOR localization to lysosomes via cholesterol. (A-B) shp16 SKMEL28 cells were transduced with lentivirus expressing shRNA targeting ATR (shATR). Where indicated, cells were supplemented with 50μM cholesterol. **(A)** Representative images of mTOR and LAMP2 immunofluorescence staining (left), which is quantified on the right. **(B)** Representative images of TSC2 and LAMP2 immunofluorescence staining (left), which is quantified on the right. Scale bar = 20μm. Representative data from one of three independent experiments is shown. Graphs represent mean ± SD. Oneway ANOVA. ****p<0.001, ns = not significant. **(C)** Schematic of the proposed model linking ATR to mTORC1 signaling via *de novo* cholesterol synthesis.

## DISCUSSION

Prior work found that ATR is upstream of mTORC1 under replication stress (Lamm et al., 2020). However, whether ATR influences mTORC1 in unperturbed cells is unknown. Moreover, mTORC1 is a central nutrient senor, and the metabolic mechanism linking ATR to mTORC1 has never been described. Here we found that ATR promotes mTORC1 activation both in unperturbed cells and those with p16 knockdown. This occurs through LSS and cholesterol and is independent of both CHK1 and the TSC complex. We validated these effects using both siRNA/shRNA-mediated knockdown of ATR, as well as with the ATR inhibitor AZD6738. Both ATR and mTOR belong to the PI3K-like kinase (PIKK) family, and previous generation ATR inhibitors have known offtarget effects on mTOR, underscoring the importance of genetic ATR knockdown in this study. An off-target effect of AZD6738 on mTOR can also be ruled out in this study by the rescue with lanosterol/cholesterol (**Fig. 3, S3**), and by the sensitization upon p16 knockdown (**Fig. S2E**). Thus, these results firmly establish a link between ATR, cholesterol signaling, and mTORC1 activity.

There is a bidirectional cross-talk between mTORC1 and DNA damage. Interestingly, most studies have shown that mTORC1 influences the DNA damage response (Danesh Pazhooh et al., 2021; Ma et al., 2018), with only a handful demonstrating that ATR or ATM are upstream of mTORC1 activation (Alexander et al., 2010; Lamm et al., 2020). One prior report found that replication stress, and therefore ATR activity, promotes nuclear mTORC1 activation (Lamm et al., 2020), although no direct mechanism was identified and whether mTORC1 can be activated outside of the lysosome remains an open question. Here we found that ATR promotes mTORC1 activation (**Fig. 1-2**), through increasing cholesterol levels and mTORC1 localization to the lysosome (**Fig. 3-4**). DNA damage has also been shown to induce or rewire lipid metabolism (Carroll et al., 2015; Hammerquist et al., 2021; Hamsanathan et al., 2022; Xiao et al., 2020), although to our knowledge no prior studies have examined cholesterol in this context. We found that ATR increases expression of LSS, a critical enzyme in *de novo* cholesterol synthesis (**Fig. 3**). How exactly ATR expression and/or activity promotes this pathway remains to be investigated. ATR was long thought of as a nuclear protein, while LSS is cytoplasmic. However, ATR has also recently been found in the cytoplasm (Biswas et al., 2022; Hilton et al., 2016; Postigo et al., 2017). Given that ATR is a kinase (Saldivar et al., 2017), it is possible ATR directly phosphorylates LSS to promotes its activity or stability. Investigating this will be the aim of future studies.

We found that ATR suppresses mTORC1 in part via reducing cholesterol levels, as cholesterol supplementation rescued the effect of ATR knockdown or inhibition on mTORC1 (**Fig. 3**). We demonstrated that this effect is through changes in mTOR localization at the lysosome, and independent of the TSC complex (**Fig. 4**). Prior work has elegantly shown how cholesterol promotes mTORC1 activity via LYCHOS at the lysosome (Shin et al., 2022). Whether LYCHOS is the mediator of the changes in mTORC1 activity downstream of ATR remains to be tested. It is also interesting to speculate whether increased cholesterol has other signaling effects beyond increasing mTORC1 activity. Cholesterol is a critical component of the plasma membrane but can also lead to increased steroid hormones, vitamin D, and bile acids (Centonze et al., 2022; Liu et al., 2023). We and others have found that p16 knockdown cells proliferate faster than controls (Buj and Aird, 2019; Fung et al., 2013), and thus likely need increased membrane synthesis to support more frequent cell division. Additionally, both steroids and bile acids promote pro-tumorigenic signaling pathways (Centonze et al., 2022; Liu et al., 2023; Tangudu et al., 2024). Bile acids also induce ROS (Orozco-Aguilar et al., 2021), which may induce and/or reinforce the DNA damage observed in p16 knockdown cells (Tangudu et al., 2024). Together, this likely indicates that cholesterol metabolism may have pleotropic effects on these cells.

In summary, we report a novel link between ATR and mTORC1 via LSS and cholesterol. These studies have a number of potential implications. In cancer, ATR inhibitors are currently under clinical development (Priya et al., 2023), and our results demonstrate that these inhibitors will have pleotropic effects outside of the DNA damage response that may influence therapeutic efficacy. It also raises the interesting possibility that previously reported effects of ATR on metabolism could be mediated through its effects on mTORC1. Moreover, since ATR signals to mTORC1 in normal cells (**Fig. 1**), our data suggest that there may be side effects of ATR inhibitors on both cholesterol synthesis and mTORC1 signaling that should be considered in clinical application of these therapeutics. Additionally, the DDR and mTORC1 are both deregulated during aging (Johnson et al., 2013; Schumacher et al., 2021). Thus, it is interesting to speculate whether cholesterol also connects these signaling pathways during aging or other pathologies associated with increased DNA damage. Finally, recent work has found that ATR plays an important role in normally proliferating cells, such as T cells (Menolfi et al., 2023; Sugitani et al., 2022). Therefore, this pathway may be of broad relevance to normal physiology.

## Acknowledgements

This work was supported by grants from the National Institutes of Health (R37CA240625 to K.M.A., R01CA259111 to K.M.A. and N.W.S, R35GM155379 to A.J.V., T32GM133332 to A.R.C., and S10OD023402 and S10OD032141 to S.L.G.), the Dept of Defense (HT9425-23-1-0288 to A.J.V.), the Melanoma Research Foundation (to R.B.G.), the Ludwig Institute for Cancer Research Princeton Branch (to A.J.V.), and the Ovarian Cancer Research Alliance (MIG-2023-2-1018 to A.U.). This project has been made possible in part by grant number 2023-329680 (to K.M.K.) from the Chan Zuckerberg Initiative DAF, an advised fund of Silicon Valley Community Foundation. Research reported in this publication was supported by the National Cancer Institute of the National Institutes of Health under Award Number P30CA047904.

## Author Contributions

**Naveen Kumar Tangudu:** Investigation, Methodology, Visualization, Writing – Review & Editing. **Alexandra N. Grumet**: Investigation, Methodology, Visualization, Writing – Review & Editing. **Richard Fang:** Investigation, Writing – Review & Editing. **Raquel Buj:** Investigation, Visualization, Writing – Review & Editing. **Aidan Cole:** Investigation, Writing – Review & Editing. **Apoorva Uboveja:** Investigation, Writing – Review & Editing. **Amandine Amalric:** Investigation, Writing – Review & Editing. **Baixue Yang:** Investigation. **Zhentai Huang:** Investigation. **Cassandra Happe:** Investigation. **Mai Sun:** Investigation. **Stacy L. Gelhaus:** Methodology, Supervision, Funding Acquisition. **Matthew L. MacDonald:** Methodology, Supervision, Funding Acquisition. **Nadine Hempel:** Writing – Review & Editing**. Nathaniel W. Snyder:** Writing – Review & Editing, Funding Acquisition. **Katarzyna M. Kedziora:** Methodology, Investigation. **Alexander J. Valvezan:** Conceptualization, Visualization, Writing – Original Draft, Writing – Review & Editing, Supervision, Project Administration, Funding Acquisition. **Katherine M. Aird:** Conceptualization, Visualization, Writing – Original Draft, Writing – Review & Editing, Supervision, Project Administration, Funding Acquisition.

## Materials and Methods

### Cell Lines

SKMEL28, RPMI-7951, and HeLa cells were purchased from ATCC. SKMEL28 cells were cultured in DMEM (Fisher Scientific cat# MT10013CV,) supplemented with 5% Fetal Bovine Serum (BioWest, cat# S1620). RPMI-7951 cells were cultured in MEM (Fisher Scientific cat#MT10009CV) supplemented with 5% Fetal Bovine Serum (BioWest, cat# S1620). HEK293 and HeLa cells were cultured in DMEM (Fisher Scientific cat# MT10013CV,) supplemented with 5% Fetal Bovine Serum (BioWest, cat# S1620). MEFs were cultured in DMEM (Fisher Scientific cat# MT10013CV,) supplemented with 5% Fetal Bovine Serum (BioWest, cat# S1620).Cells were supplemented with 1% Penicillin/Streptomycin (Fisher Scientific, cat#15-140-122). All cell lines were routinely tested for mycoplasma as described in (Uphoff and Drexler, 2005).

### Lentiviral packaging and infection

Lentiviral vectors were packaged using the ViraPower Kit (Invitrogen, cat#K497500) following the manufacturer’s instructions. Retrovirus was generated as we have previously published using QNX cells (Buj et al., 2019). Cells were infected with corresponding vectors for 16h and selected for 3 days with 1-3µg/ml puromycin. The plasmids are the following: pBABE-puro-H-RASG12V (Addgene, 39526); pBabe-puro (Addgene, 1764); pLKO.1-shp16 (TRCN0000010482); pLKO.1shGFP control (Addgene, cat#30323); pLKO.1-shATR (TCRN0000039615, TCRN0000039616); pLKO.1-shATM (TCRN0000038658); pLKO.1-shChk1 (TRCN0000000499, TRCN0000000502); pLKO.1-shRPTOR (TRCN0000039772); pLKO.1-LSS (TRCN0000045481, TRCN0000045482); pLKO.1-shCdkn2a (TRCN0000077816, TRCN0000362595).

### Cholesterol and Lanosterol treatment

Cells were seeded at an equal density on plates or coverslips. Post-lentiviral infection, cells were selected with puromycin for 24h. Cells were washed once with DMEM and starved in 0.5% lipid depleted FBS/0.75%-MβCD (Omega Scientific, cat# FB-50/Sigma Aldrich, Cat# C4555) contained DMEM for 2h. Cells were washed once with DMEM and supplemented with 50μM Cholesterol (Sigma Aldrich, Cat# C3045) or lanosterol (Sigma Aldrich, Cat# L5768)/0.1%-MβCD/0.5% lipid depleted FBS for 2h. Similarly, cells treated with ATR or LSS inhibitors were also supplemented with cholesterol/lanosterol as mentioned above.

### Immunofluorescence

Cells were seeded at an equal density on coverslips and fixed with 4% paraformaldehyde. Cells were washed four times with PBS and permeabilized with 0.2% Triton X-100 in PBS for 5min. For LSS staining, cells were blocked for 5 min with 3% BSA/PBS followed by incubation of corresponding Rabbit anti-LSS primary antibody (Sigma Aldrich, cat# HPA032060; 1:200) in 3% BSA/PBS for 1h at room temperature. Cells were washed three times with 1% Triton X-100 in PBS and incubated with secondary antibody (Cy3 Goat Anti-Rabbit (Jackson Immuno, cat# 111165-003, 1:5000), and Cy3-AffiniPure Donkey Anti-Mouse (Jackson Immuno, cat# 715-165-150, 1:5000)) in 3% BSA/PBS for 1h at room temperature. Cells were then incubated with 0.15 µg/ml DAPI for 1 min, washed three times with PBS, mounted with fluorescence mounting medium (9 ml of glycerol [BP229-1; Fisher Scientific], 1 ml of 1× PBS, and 10 mg of p-phenylenediamine [PX0730; EMD Chemicals]; pH was adjusted to 8.0–9.0 using carbonate-bicarbonate buffer [0.2 M anhydrous sodium carbonate, 0.2 M sodium bicarbonate]) and sealed. Images of LSS staining, along with images of nuclei stained with DAPI, were segmented using the pre-trained Cellpose model ‘cyto3’ (Cellpose v. 3.0.7) (Stringer et al., 2021). Objects that touched the border of the image and objects below and above set area thresholds (5k – 100k pixels) were excluded from further analysis. The fluorescence signal of LSS was measured across entire cell regions and background-corrected by subtracting the median signal of the entire image. For mTOR/LAMP2 and TSC2/LAMP2 staining, cells were blocked with Intercept Blocking Buffer (Licor #927-70001) diluted 1:1 in PBS for 1 hr at room temperature. Cells were incubated overnight at 4°C in primary antibodies: TSC2 (CST #4308, 1:1250) or mTOR (CST #2938, 1:200) and LAMP2 (Santa Cruz #sc18822, 1:100). Cells were then washed with PBS, incubated with secondary antibodies conjugated to Alexa Fluor 488 (ThermoFisher Scientific #A-11001, 1:1000) and Cy3 (Jackson ImmunoResearch #111-165-144, 1:1000) for 1 hour at room temperature, washed with PBS again, and mounted on slides with Fluoromount G (SouthernBiotech #0100-01) for imaging using a Zeiss LSM 900 confocal microscope. TSC2:LAMP2 and mTOR:LAMP2 colocalization was quantified using the Zeiss ZenBlue 3.3 analysis suite.

### Western blotting

Cells lysates were collected in 1X sample buffer (2% SDS, 10% glycerol, 0.01% bromophenol blue, 62.5mM Tris, pH 6.8, 0.1M DTT) or lysis buffer containing 20 mM Tris pH 7.5, 140 mM NaCl, 1 mM EDTA, 10% glycerol, 1% Triton X-100, 50 mM NaF, 1 mM DTT, with protease inhibitor cocktail (Sigma #P8340), phosphatase inhibitor cocktail #2 (Sigma #P5726), and #3 (Sigma #P0044) used at 1:100 each, and then boiled to 95°C for 5-10 min. Cell lysates were sonicated for 10-15sec. Protein concentration was determined using the Bradford assay (Bio-Rad, cat#5000006). An equal amount of total protein was resolved using SDS-PAGE gels and transferred to nitrocellulose membranes (Fisher Scientific) at 110mA for 2h at 4°C. Membranes were blocked with 5% nonfat milk or 4% BSA in TBS containing 0.1% Tween-20 (TBS-T) for 1h at room temperature. Membranes were incubated overnight at 4°C in primary antibodies in 4% BSA/TBS + 0.025% sodium azide. Membranes were washed 4 times in TBS-T for 5 min at room temperature after which they were incubated with HRP-conjugated secondary antibodies (Cell Signaling Technology) for 1 h at room temperature. After washing 4 times in TBS-T for 5 min at room temperature, proteins were visualized on film after incubation with SuperSignal West Pico PLUS Chemiluminescent Substrate (ThermoFisher, Waltham, MA). Primary antibodies: Rabbit anti-p16 INK4A (Cell Signaling Technology, 29271; 1:1000); Phospho-Chk1 (Ser345) (Cell Signaling Technology, Cat# 2348, 1:1000); Mouse anti Chk1 (Cell Signaling Technology, Cat# 2360, 1:1000); Rabbit anti-Phospho-Chk2 (Thr68) (Cell Signaling Technology, Cat# 2197, 1:1000); Rabbit anti-Chk2 (Cell Signaling Technology, Cat# 2662, 1:1000); Rabbit anti-LSS (Sigma Aldrich, cat# HPA032060; 1:1000); mouse anti-β-actin (Sigma Aldrich, cat# A1978; 1:10,000); Mouse anti-vinculin (Sigma Aldrich, cat# V9131; 1:10,000); Rabbit anti-ATR (Bethyl, Cat # A300-138A, 1:1000); Rabbit antiphopsho-p70 S6 Kinase (T389) (Cell Signaling Technology, Cat # 9234, 1:1000); Rabbit antip70 S6 Kinase (Cell Signaling Technology, Cat # 2708, 1:1000); Secondary antibodies: Anti-mouse IgG, HRP-linked (Cell Signaling Technology, 7076; 1:10,000 and 1:5000), AntiRabbit IgG, HRP-linked (Cell Signaling Technology, 7074; 1:5000).

### Analysis of The Cancer Genome Atlas patient data

RPPA Data were extracted from The Cancer Genome Atlas (TCGA) Skin Cutaneous Melanoma Firehose Legacy using cBioportal. Patients were divided based on homozygous deletion (or not) of *CDKN2A*.

### Proteomics

SKMEL28 cells were homogenized in 50mM TEAB, 5% SDS. Total protein was measured by microBCA (Pierce). 500 µg total protein was digested on S-Trap Midi’s (Protifi) per manufacture protocol and desalted on Peptide Desalting Spin Columns (Pierce). Phosphopeptides were enriched on an AssayMAP Bravo (Agilent) with Fe3+ column. LC-TIMS-MS/MS analysis was carried out using a nanoElute UHPLC system (Bruker Daltonics, Bremen, Germany) coupled to the timsTOF Pro mass spectrometer mass spectrometer (Bruker Daltonics), using a CaptiveSpray nanoelectrospray (Bruker Daltonics). Roughly 100 ng of peptide digest or phosphopeptide enrichment was loaded on a capillary C18 column (25 cm length, 75 μm inner diameter, 1.6 μm particle size, 120 Å pore size; IonOpticks, Fitzroy, VIC, AUS). Peptides were separated at 55°C using a 60 min gradient at a flow rate of 300 nL/min (mobile phase A (MPA): 0.1% FA; mobile phase B (MPB): 0.1% FA in acetonitrile). A linear gradient of 2–35% MPB was applied for 60 min, followed by a 5 min wash at 95% MFB before equilibrating the column at 2% MFB for 6 min. The timsTOF Pro was operated in PASEF mode collecting full scan mass spectra from 100 and 1700 *m*/*z*. Ion mobility resolution was set to 0.60–1.60 V·s/cm over a ramp time of 100 ms. Data-dependent acquisition was performed using 10 PASEF MS/MS scans per 1.1 second cycle. Active exclusion time window was set to 0.4 min, and the intensity threshold for MS/MS fragmentation was set to 2.5e4 while low *m*/*z* and singly charged ions were excluded from PASEF precursor selection. MS/MS spectra were acquired via ramped collision energy as function of ion mobility.

The nLC-MS/MS data were analyzed with the MaxQuant software suite (version 2.1.3) (Cox and Mann, 2008). The Andromeda protein identification search engine (Cox et al., 2011) and a SwissProt human protein database (downloaded on November 11, 2022 with 20,403 entries) was utilized with default settings for Orbitrap instruments. The parameters used included a precursor mass tolerance of 20 ppm for the first search and 4.5 ppm for the main search, a product ion mass tolerance of 0.2 Da, and a minimum peptide length of 5 amino acids. Trypsin was set as the proteolytic enzyme with a maximum of two missed cleavages allowed. The enzyme specifically cleaves peptide bonds C-terminal of arginine and lysine if they are not followed by proline. Carbamidomethylation of cysteine was set as a fixed modification. Oxidation of methionine, deamination of both asparagine and glutamine, and acetylation of the protein N-terminus were set as variable modification. Phosphorylation of serine, threonine, and tyrosine was set as a variable modification for phosphopeptides. A 1% false discovery rate (FDR) was used to filter the peptide identification results. The integrated feature intensities provide a relative measure of abundance for each feature at the peptide level and used in all subsequent analyses. Proteomics data can be found in **Table S1**.

### Filipin staining and analysis

Cells were cultured on coverslips and fixed with 4% paraformaldehyde. Cells were incubated with 0.1mg/ml filipin III (Cayman Chemical, cat# 70440) for 45-60min. Coverslips were washed twice with PBS, mounted, and sealed. Images of Filipin staining were segmented using the pre-trained Cellpose model ‘cyto3’ (Cellpose v. 3.0.7) (Stringer et al., 2021). Objects that touched the border of the image and objects below and above set area thresholds (25k – 100k pixels) were excluded from further analysis. The fluorescence signal of Filipin was measured across entire cell regions with flat background correction (200 a.u.).

### Quantification and Statistical Analysis

GraphPad Prism (version 10.0) was used to perform statistical analysis. Point estimates with standard deviations are reported, and the appropriate statistical test was performed using all observed experimental data. All statistical tests performed were two-sided and p-values < 0.05 were considered statistically significant.

### Data Availability Statement

All data generated or analyzed during this study are included in this published article and its supplementary information files.

## SUPPLEMENTAL FIGURES

**Supplemental Figure 1.**
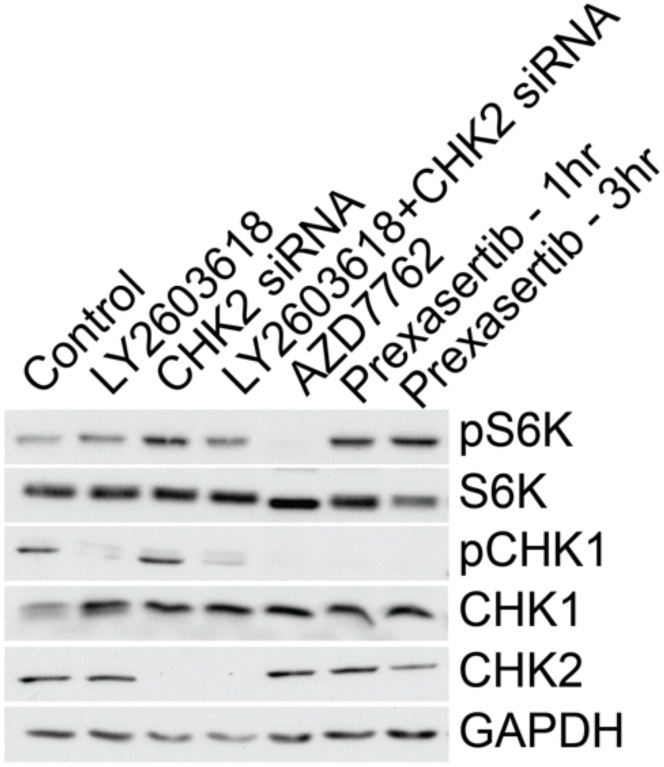
S**u**ppression **of CHK1 and CHK2 do not markedly decrease mTORC1 activity. Related to Figure 1**. HeLa cells were treated with the CHK1 inhibitor LY2603618 (2µM, 3 hrs), CHK2 siRNA (50 nM, 48 hrs), the dual CHK1/CHK2 inhibitor AZD7762 (2µM, 3 hrs), or the dual CHK1/CHK2 inhibitor Prexasertib [100 nM, 1 (lane 6) or 3 (lane 7) hrs], and the indicated proteins were assessed by western blotting. GAPDH was used as a loading control. All western blots are representative data from at least 3 independent experiments.

**Supplemental Figure 2.**
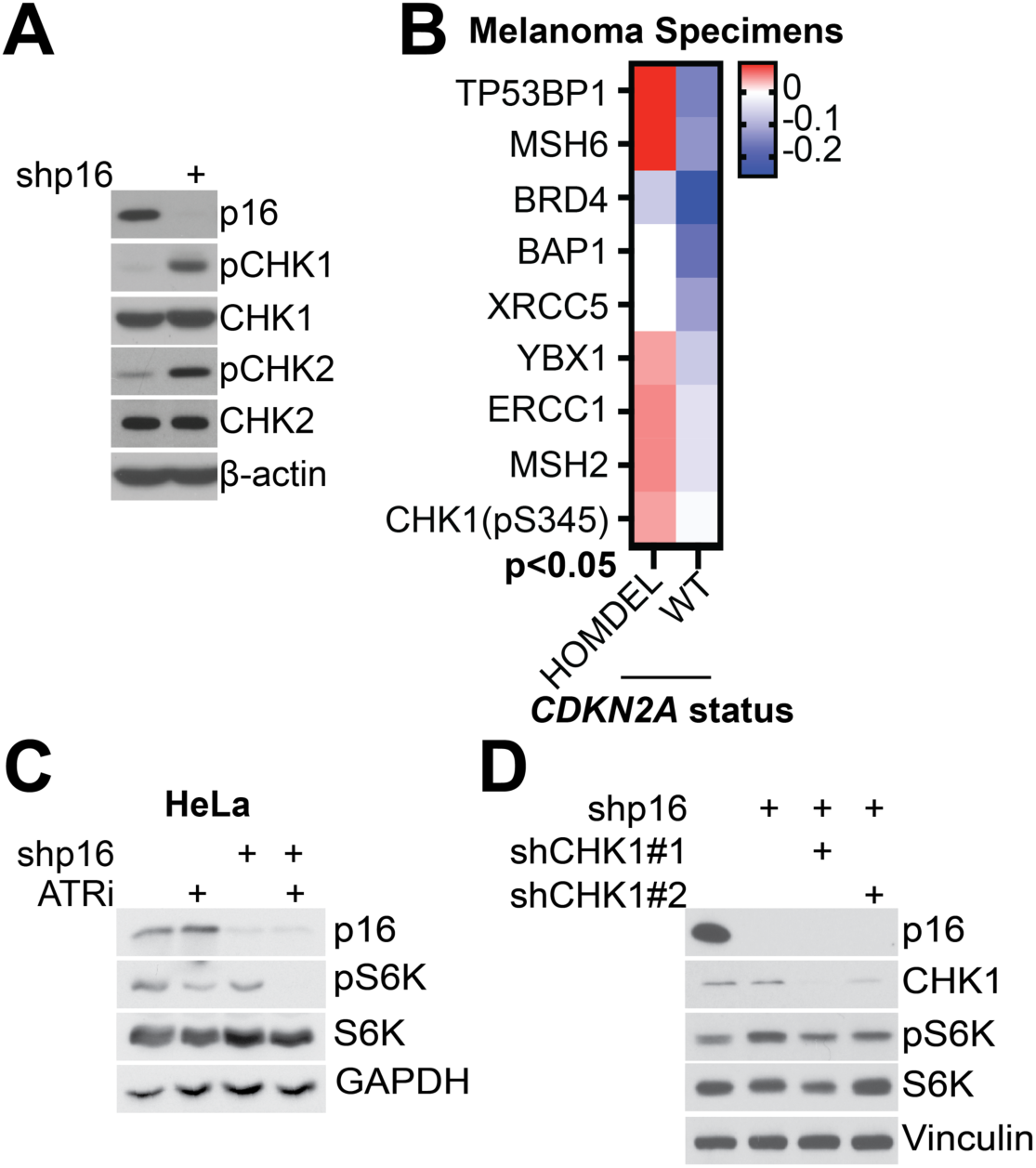
p**1**6 **knockdown increases activation of the ATR/ATM pathways; ATR-mediated mTORC1 activation in p16 knockdown cells is independent of CHK1. Related to Figure 2. (A)** SKMEL28 cells were transduced with a lentivirus expressing a short hairpin targeting GFP as a control or p16 (shp16), and the indicated proteins were assessed by western blotting. β-actin was used as loading controls. **(B)** RPPA results from Melanoma patient samples show upregulation of proteins related to the DNA damage response and repair in tumors with homozygous deletion (HOMDEL) of *CDKN2A* (encoding p16) compared to wildtype (WT) *CDKN2A* (all proteins p<0.05). Graphs represent mean ± SD. T-test. *p<0.05. **(C)** HeLa cells were transduced with a lentivirus expressing a short hairpin targeting GFP as a control or p16 (shp16) and treated with 0.5µM AZD6738 (ATRi) for 30 min. The indicated proteins were assessed by western blotting. GAPDH was used as a loading control. **(D)** Same as **(A)**, but shp16 cells were transduced with a lentivirus expressing a short hairpin targeting GFP or two short hairpins targeting CHK1 (shCHK1 #1 and #2), and the indicated proteins were assessed by western blotting. Vinculin was used as a loading control. Representative data from three independent experiments is shown for all western blots.

**Supplemental Figure 3.**
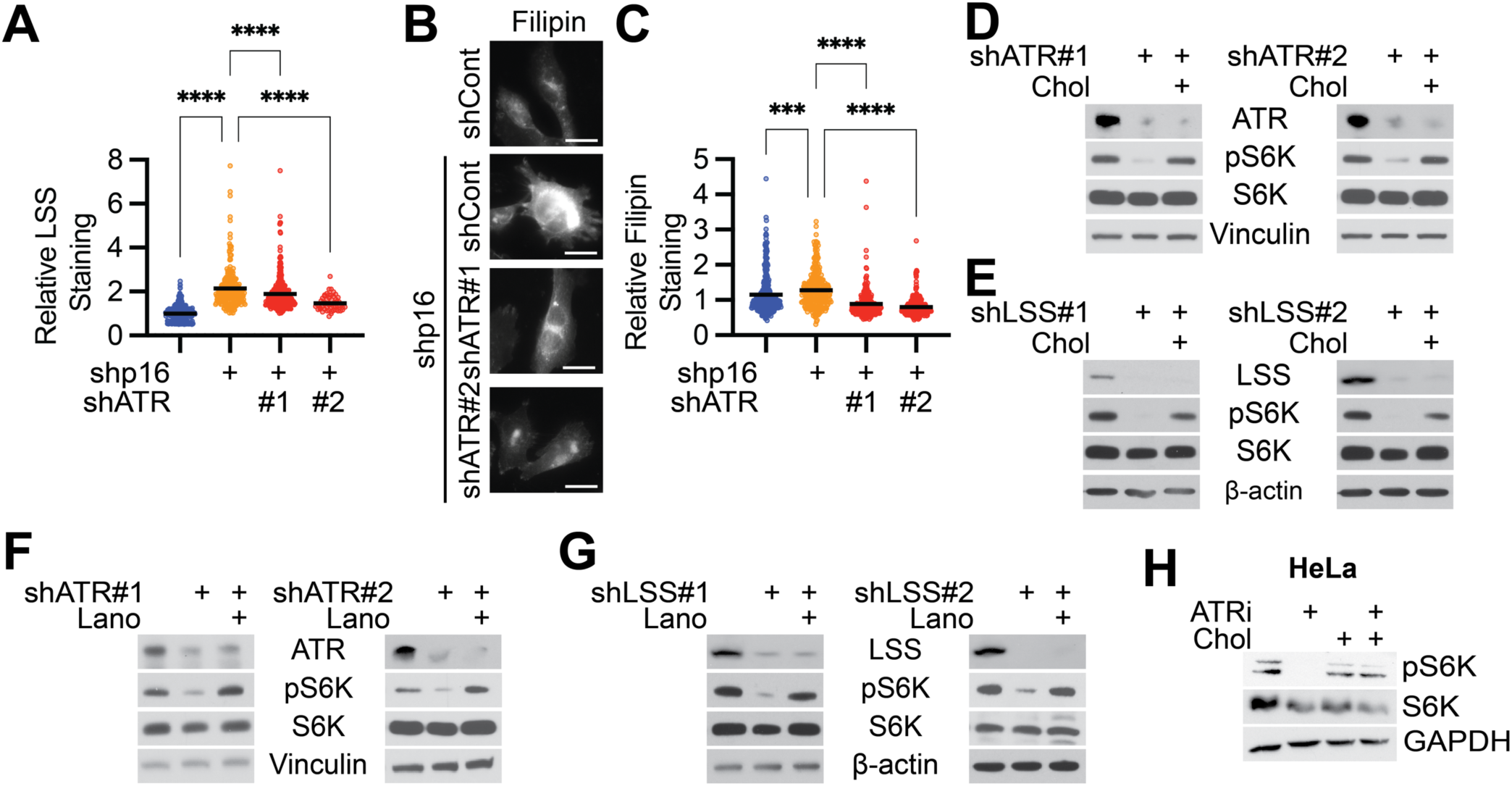
L**S**S **and cholesterol are downstream of ATR. Related to Figure 3. (A-B)** RPMI-7951 cells were transduced with lentivirus expressing shRNA targeting GFP as a control or p16 (shp16) with or without lentivirus expressing shRNA targeting ATR (shATR #1 and #2). **(A)** LSS expression was assessed by immunofluorescence staining and quantified. **(B-C)** Cholesterol abundance was assessed by filipin staining **(B)** and quantified **(C)**. Scale bar = 20μm. **(D-E)** shp16 RPMI-7951 cells were transduced with lentivirus expressing targeting ATR (shATR #1 and #2) **(D, F)** or targeting LSS (shLSS #1 and #2) **(E, G)** and supplemented with 50μM cholesterol or 50μM lanosterol where indicated. (**H**) shp16 HeLa cells were treated with vehicle or 0.5µM AZD6738 (ATRi) for 30 min in the presence or absence of supplementation with 50μM cholesterol. Representative data from one of three independent experiments is shown for all panels. Graphs represent individual normalized values and mean. One-way ANOVA. ***p<0.005, ****p<0.001.

**Supplemental Figure 4.**
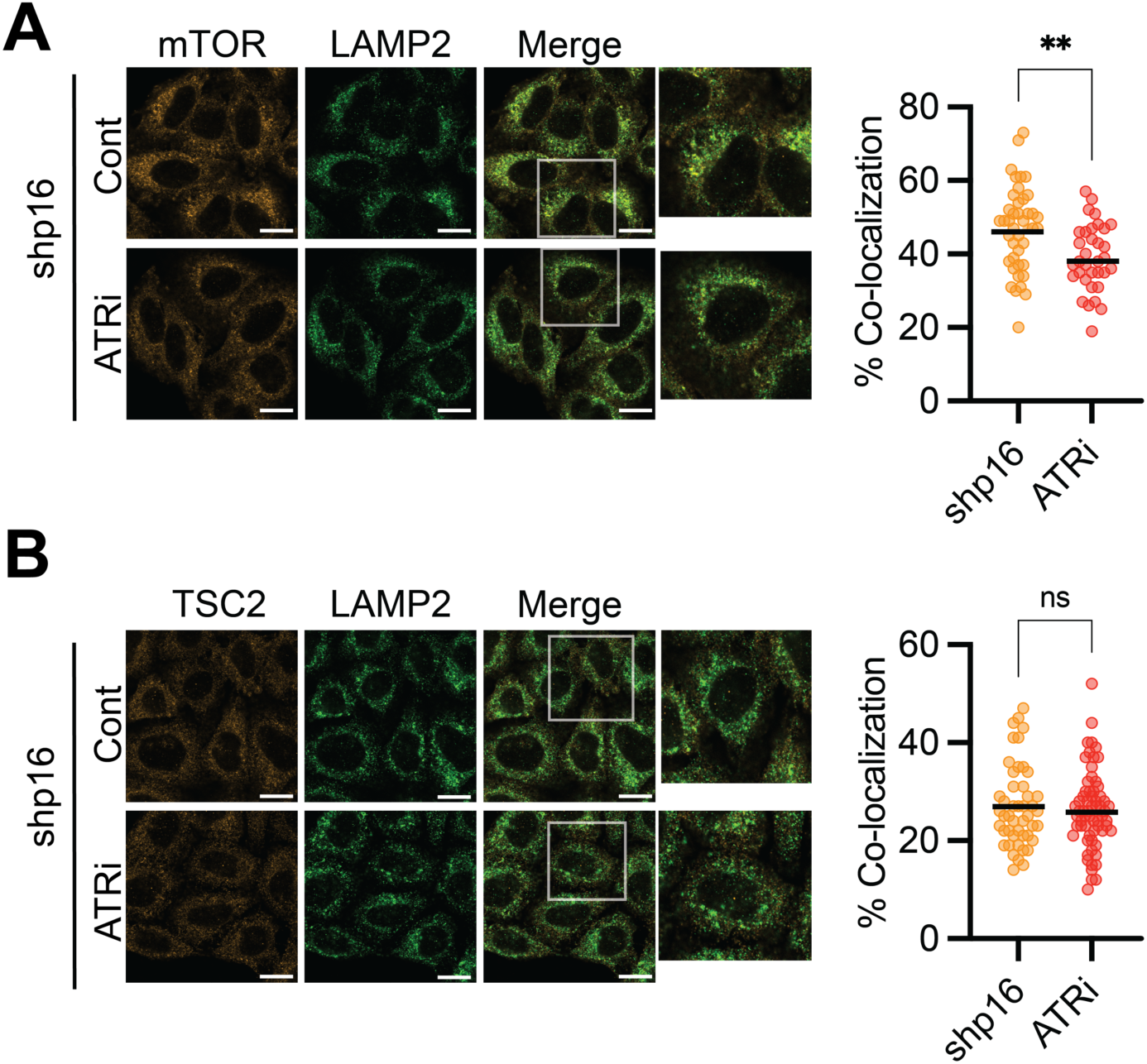
A**T**R **inhibition decreases mTOR at the lysosome but has no effect on TSC2. Related to Figure 4. (A-B)** shp16 HeLa cells treated for 30 min with 0.5μM AZD6738 ATR inhibitor (ATRi) or vehicle. **(A)** Representative images of mTOR and LAMP2 immunofluorescence staining (left), which is quantified on the right. **(B)** Representative images of TSC2 and LAMP2 immunofluorescence staining (left), which is quantified on the right. Scale bar = 20μm. Representative data from one of three independent experiments is shown. Graphs represent individual normalized values and mean. Student’s t-test. **p<0.0015, ns = not significant.

